# An output-null signature of inertial load in motor cortex

**DOI:** 10.1101/2023.11.06.565869

**Authors:** Eric A. Kirk, Keenan T. Hope, Samuel J. Sober, Britton A. Sauerbrei

**Affiliations:** Case Western Reserve University School of Medicine, Department of Neurosciences; Emory University, Department of Biology

## Abstract

Coordinated movement requires the nervous system to continuously compensate for changes in mechanical load across different contexts. For voluntary movements like reaching, the motor cortex is a critical hub that generates commands to move the limbs and counteract loads. How does cortex contribute to load compensation when rhythmic movements are clocked by a spinal pattern generator? Here, we address this question by manipulating the mass of the forelimb in unrestrained mice during locomotion. While load produces changes in motor output that are robust to inactivation of motor cortex, it also induces a profound shift in cortical dynamics, which is minimally affected by cerebellar perturbation and significantly larger than the response in the spinal motoneuron population. This latent representation may enable motor cortex to generate appropriate commands when a voluntary movement must be integrated with an ongoing, spinally-generated rhythm.

## 1. Introduction

The ability to perform the same movement repeatedly in a changing environment is a hallmark of skilled motor control. Inertial load is a key environmental variable which changes with the distribution of mass across the body and must be countered with appropriately-scaled motor commands. For example, raising a coffee cup to the lips when the cup is empty and full requires different patterns of muscle activity. Similarly, the motor output generated during walking in bare feet must be adjusted when heavy boots and a backpack are worn. Such adjustments pose a demanding challenge for neural control, which is distributed across multiple interacting systems, including the motor cortex, cerebellum, brainstem, spinal cord, and muscle receptors (Fig. 1A).

**FIGURE 1.**
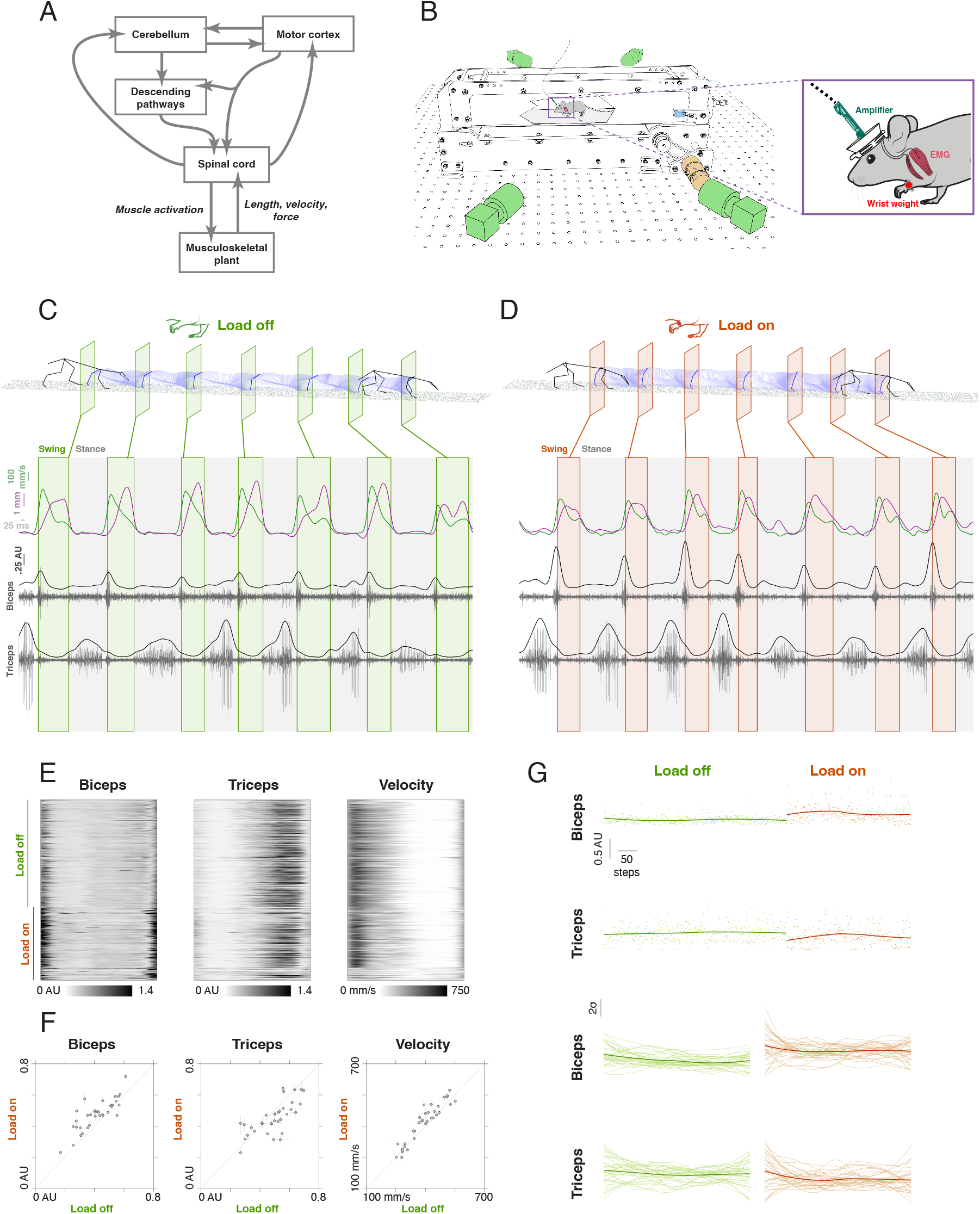
An adaptive locomotion task in freely-moving mice. (A) Block diagram illustrating key circuits involved in adaptation to mechanical loads. (B) Experimental rig. Mice were trained to trot on a motorized treadmill at 20 cm/s. Behavior was captured with four synchronized cameras, and electromyograms (EMG) recorded in the biceps brachii and triceps brachii muscles. (C) Kinematics and EMG during locomotion without a load. Upper: 3D pose estimates, with swing onset indicated in green rectangles. Lower: upward position (magenta) and forward velocity (green) of fingertip, and raw and smoothed biceps and triceps EMG (gray). (D) Kinematics and EMG during locomotion with a 0.5 g load attached to the wrist. (E) Smoothed, step-aligned biceps EMG, triceps EMG, and forward finger velocity over one experimental session. (F) Median biceps activity, triceps activity, and forward finger velocity for all sessions, load-on vs. load-off (n = 34 sessions, n = 7 mice). Lines indicate bootstrapped 95% confidence intervals. (G) Time course of EMG amplitude changes. Upper: biceps and triceps amplitudes across a single session. Points correspond to individual steps, and lines indicate a loess estimate of the trend. Lower: loess estimates for all sessions. To enable comparison across sessions, each curve was z-scored and stretched to unit duration.

In the context of voluntary movement, studies in nonhuman primates have demonstrated that the motor cortex drives the generation of forces to move the upper limb and to compensate for loads^1,2^. Ablation of the pyramidal tract causes deficits in manual dexterity^3^ and the time course of force development^4^, and cortical firing rates are strongly modulated by force magnitude and direction during upper limb movements^5–7^ and isometric contractions^8^. In reaching, cortical neurons are sensitive to the direction and magnitude of loads during posture and movement^5^, and their responses can shift substantially between these two contexts^9^. Furthermore, several observations suggest that load-related responses in motor cortex might be driven, in part, by ascending cerebellar input. Cooling of the cerebellar dentate nucleus attenuates long-latency motor cortical responses to impulse torques during voluntary elbow movement^10,11^, though activity during holding against a load is minimally affected by this manipulation^12^, and disruption of cerebellar output with high-frequency electrical stimulation can partially suppress cortical activity in an isometric wrist task^13^.

The complexity and heterogeneity of the motor cortical population pose a significant challenge to understanding its role in control^14,15^. For example, a neuron’s response to load during reaching cannot be accurately predicted from its load sensitivity during posture^9^, and directional tuning can change substantially between movement preparation and execution^16^ and throughout a reach^14^. A powerful emerging approach to this complexity focuses less on the information represented by individual neurons and more on the coordinated dynamics across the cortical population, how these dynamics are related to features of the task, and how they are generated by interactions across brain areas^17–20^. This approach has helped explain several perplexing features of cortical activity, such as the observation that large changes in firing rate can occur during motor preparation without evoking movement. As a movement is planned, cortical activity changes in directions, termed output-null dimensions, along which the net effect of cortical output on muscle activity is constant^21,22^. These changes enable the cortical population state to be set to the appropriate initial condition from which dynamics can evolve during movement execution.

Given the central role of motor cortical dynamics in voluntary limb movements, how might these dynamics contribute to load compensation in rhythmic movements which are coordinated by an intrinsic spinal network? In mammalian overground locomotion, a spinal central pattern generator (CPG) governs the basic pattern of flexor-extensor and left-right limb alternation, can operate independently of the brain and sensory feedback^23–25^, and is controlled by networks in the midbrain and brainstem that determine locomotor initiation and speed^26–28^. Motor cortex is not necessary for locomotion over a flat surface, but is required when precision demands are increased during steps over obstacles or across a horizontal ladder^29–33^. Some adjustments for mechanical load are implemented by subcortical structures: in walking premammilary cats, for instance, loading of an ankle extensor tendon increases the activation of the corresponding muscle during stance, and can suppress the CPG when large forces are applied^34^. Nonetheless, the rhythmic, step-entrained activity of some cortical cells, including pyramidal tract neurons projecting to the spinal cord and brainstem, can be modulated by speed and by loading of the limb^35,36^, suggesting that descending cortical signals may be important for the regulation of force during locomotion.

The present study aims to address three central questions. First, does motor cortex drive compensation for changes in inertial load imposed on the limbs during locomotion, as it does in voluntary movement, or is this compensation instead implemented by subcortical structures? Second, how are such loads represented in motor cortical population activity, and does the representation depend on cerebellar input? Finally, how are cortical dynamics related to the output of the nervous system at the level of the spinal motoneuron population? We address these questions in unrestrained, chronically-instrumented mice performing an adaptive locomotion task in which they must adjust motor output to compensate for a weight on the wrist. Our approach combines three-dimensional kinematic pose estimation, recordings from forelimb muscles, the motor cortex, and spinally-innervated motor units, optogenetic perturbations, and computational approaches for modeling neural population data. We find that, although inactivation of motor cortex does not attenuate load compensation, the dominant component of cortical population activity is a tonic shift imposed by the load, and is robust to optogenetic perturbation of the cerebellum. Furthermore, the geometric properties of cortical population activity in the task contrast strongly with those of the spinal motoneuron population. While cortical activity is significantly modulated by load, cerebellar perturbation, and animal speed, with cortical trajectories that maintain relatively low tangling across experimental conditions, consistent with noise-robust dynamics, the spinal motoneuron population is instead dominated by condition-invariant signals related to flexor-extensor alternation, and also exhibits higher trajectory tangling. We conclude that load-related dynamics in motor cortex do not directly drive motor compensation during locomotion, but instead constitute a latent representation of changes to the limb mechanics, which may modulate cortical commands during voluntary gait modification or alter the gain of spinal reflexes to correct for unexpected perturbations.

## 2. Results

### 2.1. Adaptation of locomotor output to changes in inertial load

Unrestrained mice were trained to trot at approximately 20 cm/s on a motorized treadmill as their movements were captured with four synchronized high-speed cameras (Fig. 1B). Three-dimensional limb kinematics were measured from video using an automatic pose estimation pipeline^37,38^, enabling extraction of fingertip position and velocity (Fig. 1C, lower: magenta and green traces) and segmentation of the session into swing and stance epochs (Fig. 1C, lower: green boxes). Electromyograms (EMG) were recorded from forelimb flexor (biceps brachii) and extensor (triceps brachii) muscles, rectified, and smoothed (Fig. 1C, lower: gray traces). At the beginning of each session, animals ran freely for 5-20 minutes. We then imposed an inertial load on one forelimb by attaching a 0.5 g weight to the wrist, increasing the moment of inertia of the radius-ulna about the elbow, and the animals ran for a second epoch of 5-10 minutes. This load, which increased the total mass of the forelimb by approximately 50%, induced a compensatory increase in elbow flexor muscle activity during swing and a corresponding suppression of extensor activity during stance (Fig. 1D). The compensation was consistent across step cycles (Fig. 1E) and sessions (Fig. 1F; signed rank test, p = 8.4e-6 for biceps, p = 7.1e-3 for triceps). Finger velocity was, on average, slightly higher in the loaded condition (Fig. 1F; p = 5.7e-4), consistent with modest overcompensation for the load. Furthermore, in contrast with adaptation to a split-belt treadmill, which unfolds over many successive steps and requires cerebellum^39,40^, this adaptation appeared to occur almost instantaneously after the load was applied (Fig. 1G).

### 2.2. Load compensation is robust to perturbation of motor cortex and cerebellum

Adjustment of motor output in different tasks requires distinct contributions from motor cortex^3,36^, cerebellum^41^, and cerebellar inputs to cortex^11^. To determine whether the observed compensation for inertial load requires motor cortex and cerebellum, we used an optogenetic approach to transiently inactivate each brain area during the task. Motor cortical perturbation experiments were performed in VGAT-ChR2-EYFP mice, which express the light-gated ion channel ChR2 selectively in inhibitory interneurons, enabling robust suppression of cortical output following illumination of the brain surface with blue light^20,42^. An optical fiber was implanted over the forelimb motor cortex (Fig. 2A, left), and animals performed treadmill locomotion with and without a 0.5 g weight on the contralateral forelimb as laser stimulation was delivered intermittently to suppress motor cortical activity (473 nm, 40 Hz, 1 s stimulus duration, randomized 1-5 s delay between stimuli). While the load induced an increase in elbow flexor muscle activity during swing and a decrease in extensor activity during stance, cortical perturbation did not attenuate this compensation (Fig. 2A, center and right). We next tested the effects of cerebellar perturbation on motor output by implanting a fiber over the forelimb area of the pars intermedia ipsilateral to the loaded forelimb in L7Cre-2 x Ai32 mice (Fig. 2B, left), which express ChR2 selectively in Purkinje cells and allow suppression of cerebellar output during laser stimulation^43^. Cerebellar perturbation did not erase the adaptation of motor output to the load; on the contrary, it produced a modest flexor muscle enhancement and extensor attenuation (Fig. 2B, center and right; Supplementary Fig. 1C-D). To quantify the effects of load, optogenetic perturbation, and speed on motor output, we fit a linear model for each experimental session and examined the distribution of the resulting coefficients (Fig. 2C; Supplemental Fig. 1C-D; see Methods). Load had a significant positive effect on elbow flexor EMG and a negative effect on extensor EMG (sign rank test, q < .05), and step frequency had positive effects on both flexor and extensor EMG. The interaction terms between load and both optogenetic perturbations were centered at zero, indicating that these perturbations failed to erase the adaptation of motor output to changes in load. Overall, these results show that load compensation in the task does not require normal motor cortical or cerebellar output.

**FIGURE 2.**
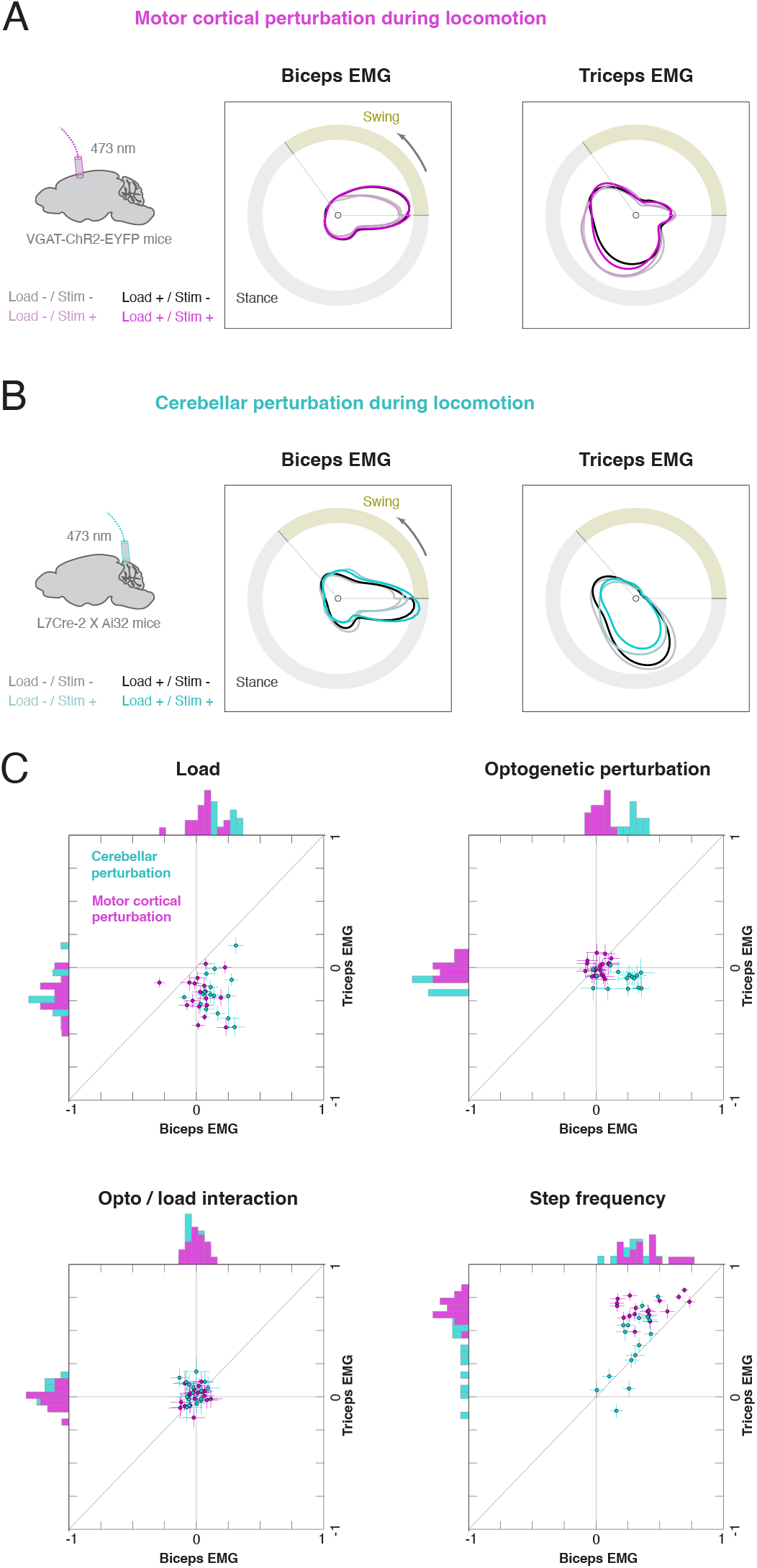
Adaptation to load is robust to cortical and cerebellar perturbation. (A) Effect of load and motor cortical perturbation on biceps and triceps EMG in a single session. Each contour corresponds to the average step-locked EMG in one of four load and optogenetic perturbation conditions. The angle represents the phase of the step cycle, and the radius the EMG magnitude at the corresponding step phase. (B) Effect of load and cerebellar perturbation on biceps and triceps EMG in a single session. (C) Regression coefficients estimating the effects of load, optogenetic perturbation, the interaction between load and optogenetic perturbation, and step frequency on biceps and triceps EMG (motor cortical perturbation in VGAT-ChR2-EYFP animals: n = 4 mice, n = 18 sessions; cerebellar perturbation in L7Cre-2 x Ai32 animals: n = 3 mice, n = 16 sessions). Each point corresponds to a single experimental session; lines denote 95% confidence intervals.

### 2.3. Load and cerebellar perturbation modulate motor cortical activity

The finding that muscle activity was unaffected by cortical inactivation suggested that the cortical dynamics in the load compensation task were output-null. The possibility remained, however, that motor cortex could still detect changes in inertial load. To measure the effects of load during locomotion on motor cortical dynamics, and to assess the dependence of these effects on cerebellar inputs, we chronically implanted high-density silicon probes in the motor cortex of L7Cre-2 x Ai32 mice, along with an optical fiber over the contralateral forelimb area of the cerebellar pars intermedia (Fig. 3A). Mice then performed the adaptive locomotion task as we recorded limb kinematics and cortical spiking (n = 710 neurons, n = 2 mice) while intermittently perturbing the cerebellum by stimulating Purkinje cells (473 nm, 40 Hz, 1 s stimulus duration, randomized 1-5 s delay between stimuli). Most neurons were synchronized with the locomotor rhythm (n = 618/710, 87.0%, q < .05, Rayleigh test with Benjamini-Hochberg correction for multiple comparisons), consistent with studies in cats^29,36^ and primates^44,45^. The effects of load and cerebellar perturbation were highly diverse across neurons. Firing rates for some cells were modulated by load (neurons 1-3, Fig. 3B), by Purkinje cell stimulation (neuron 5), or by both (neuron 6), while effects were relatively modest for others (neuron 4). Overall, 47.7% of neurons exhibited changes related to load, 24.1% to Purkinje cell stimulation, and 10.6% to both, while an interaction between load and Purkinje cell stimulation occurred in only 2.7% of cells (multi-way ANOVA for each neuron, q < .05). Among the load-sensitive neurons, 46.0% had higher firing rates in the load-on condition; among the neurons sensitive to Purkinje cell stimulation, 71.9% had a response of higher firing rates (Fig. 3C-D).

**FIGURE 3.**
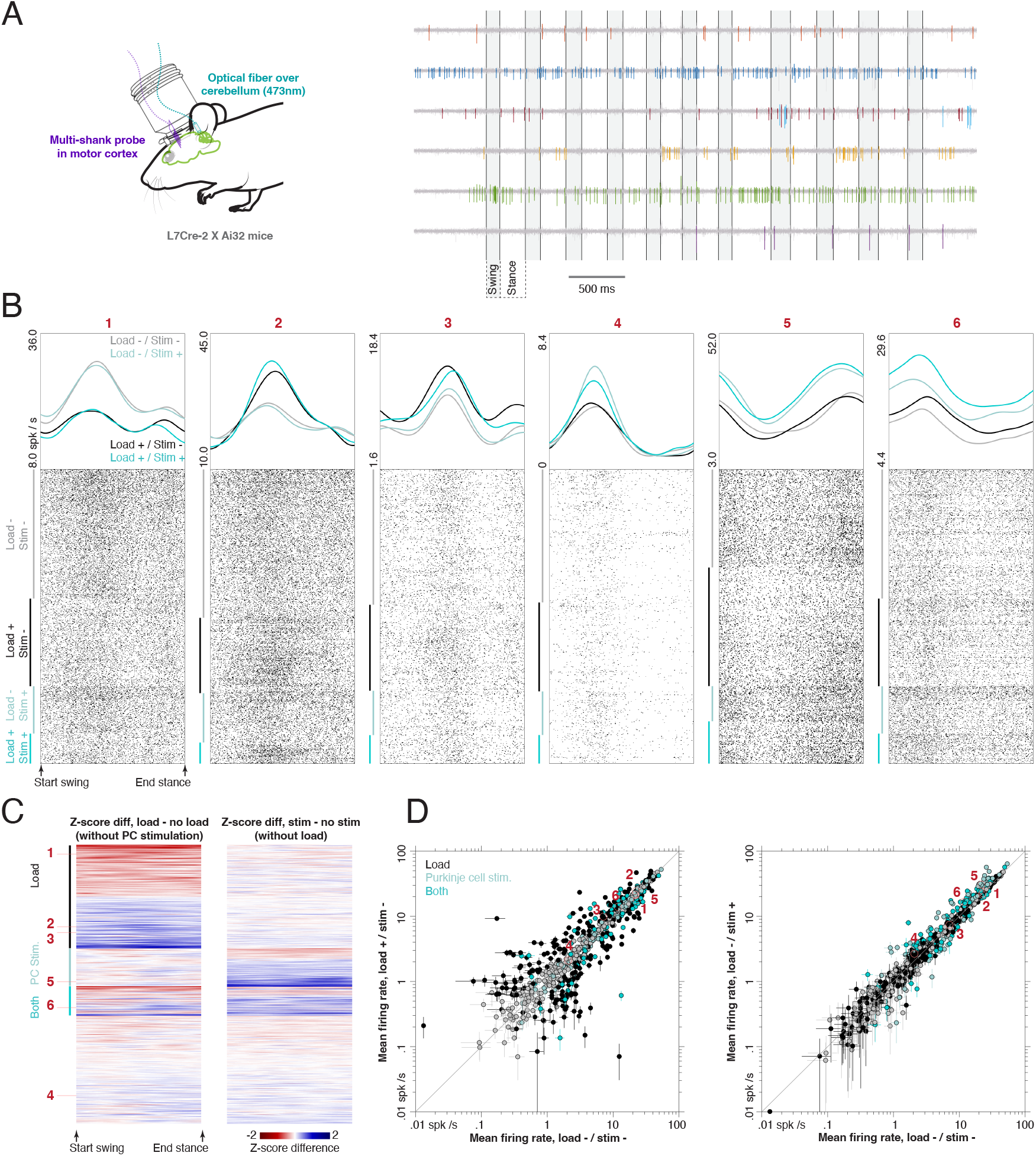
Effects of load and cerebellar perturbation on motor cortical activity. (A) Left: mice (n = 2, L7Cre-2 x Ai32) were chronically implanted with silicon probes in motor cortex and an optical fiber over the contralateral cerebellar cortex for stimulation of Purkinje cells. Right: raw data from motor cortex during locomotion. (B) Firing rates and spike rasters for six motor cortical neurons in the adaptive locomotion task. (C) Effect of load and cerebellar perturbation on motor cortical neurons (n = 710). Each row corresponds to a single neuron, and displays the difference in z-scored firing rate between load on and off conditions (left) and between Purkinje cell stimulation on and off (right). Neurons are grouped based on the detection of an effect of load (black bar), stimulation (light blue bar), both (dark blue bar), or neither (remaining neurons). (D) Mean firing rates for all neurons in load on and off conditions (left) and Purkinje cell stimulation on and off (right). Color code reflects the detection of effects of load, stimulation, or both, as in (C). Bars indicate 95% confidence intervals for the mean.

### 2.4. Cortical population dynamics in adaptive locomotion

Because the effects of load and cerebellar perturbation were heterogeneous across the sample of cortical cells, we next aimed to identify the coordinated, low-dimensional dynamics across the population. To extract a low-dimensional representation of cortical population dynamics in interpretable, task-relevant coordinates, we used demixed principal component analysis (dPCA; see Methods), which decomposes neural activity into dimensions related to specific experimental parameters while capturing most of the variance in the original firing rates^46^. For each cortical neuron, the average step-aligned firing rate was measured in twenty conditions: load on / off (two levels) x Purkinje cell stimulation on / off (two levels) x animal speed (five levels). Next, we used dPCA to find a decoder matrix that mapped the firing rates for all neurons onto a 20-dimensional latent variable space. This model explained 92.9% of the total firing rate variance and yielded scores parameterized by step phase for each dimension and condition (Fig. 4A-C), and an encoder matrix that reconstructs the measured firing rates from these scores. We observed condition-invariant signals that were modulated by step phase, but did not differ strongly across experimental parameters (Fig. 4A, X-Y axes; Fig. 4B, first column). The first condition-invariant dimension was roughly sinusoidal, with a period of one stride and a peak near the swing-stance transition. The second was qualitatively similar except for a phase shift, with a peak in mid swing, while the third was smaller in amplitude and had a period of one half stride. Taken together, the condition-invariant components accounted for 28.8% of the explained variance in cortical firing rates. Animal speed had a moderate effect on cortical dynamics (18.9% of the variance), but this was distributed broadly across multiple dimensions (Fig. 4B, fourth column; Fig, 4C). The largest speed component consisted of tonic shifts in activity, with little dependence on step phase (Fig 4B, fourth column). Dynamics in this dimension and the top two condition-invariant dimensions therefore yielded stacked elliptical trajectories that translated continuously with movement speed (Fig. 4A, right), reminiscent of motor cortical dynamics in primates performing a rhythmic cycling task^47^. Additional speed components exhibited more complex patterns, including phase and amplitude shifts in the second component.

**FIGURE 4.**
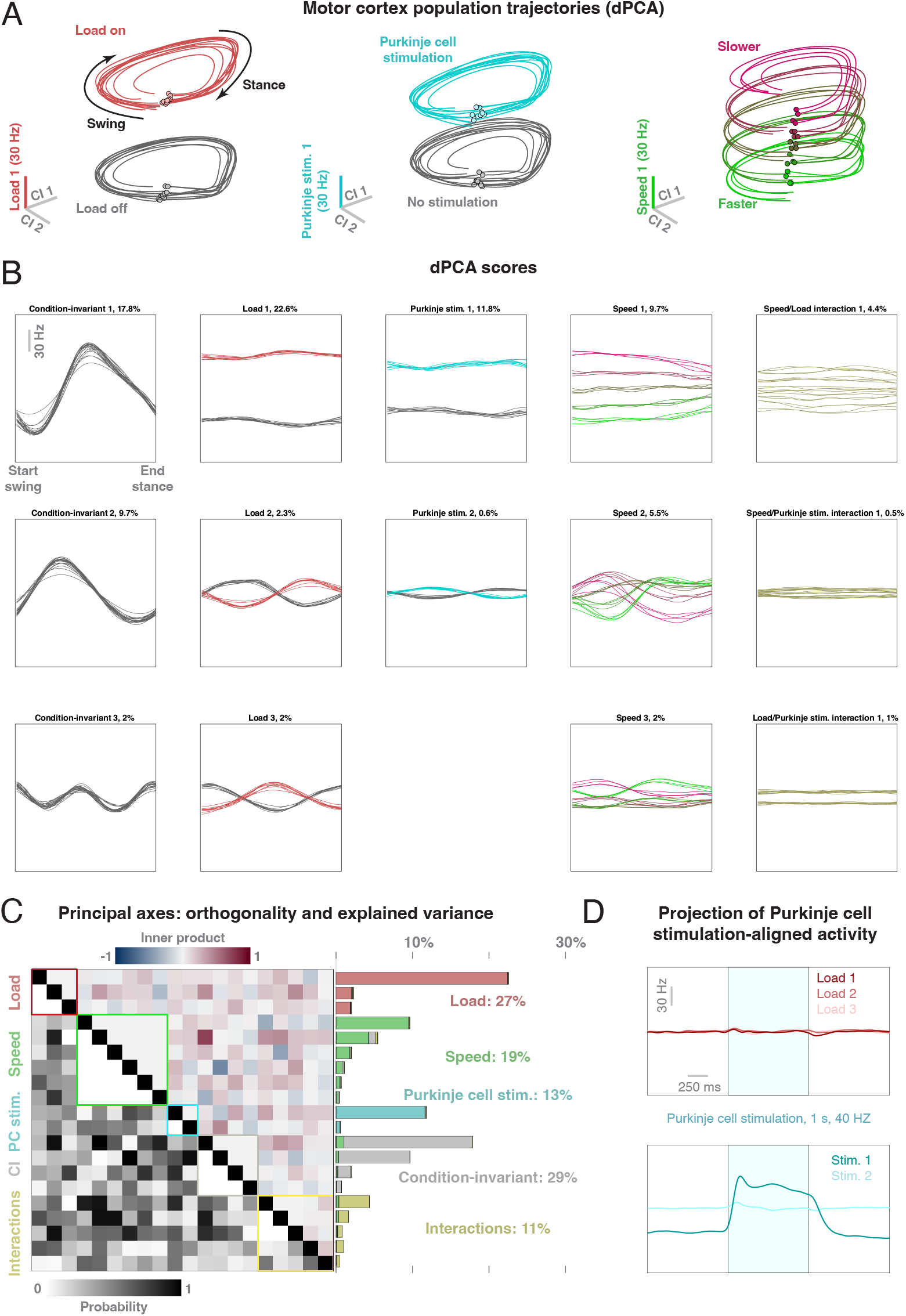
Neural population dynamics in motor cortex. (A) Step-aligned neural trajectories obtained by demixed principal component analysis (dPCA). The x and y axes in each panel represent population activity in the first two condition-invariant (CI) dimensions across twenty conditions. The z axes represent the first load dimension (left), Purkinje cell stimulation dimension (center), and speed dimension (right). (B) Step-aligned neural activity across CI, load, Purkinje cell stimulation, speed, and interaction dimensions. (C) Inner product between the principal axes (left, upper triangular), probability the inner product is larger than the observed value for randomly-oriented vectors (left, lower triangular), and fraction of variance explained by each dimension (right). (D) Projection of firing rates aligned to Purkinje cell stimulation onto load dimensions (upper) and Purkinje cell stimulation dimensions (lower).

The largest single component of cortical activity, however, was related almost purely to inertial load (Fig. 4A, left; Fig. 4B, second column, first row), accounting for 22.6% of explained variance in firing rate. This component depended only weakly on step phase, and consisted of a tonic shift in activity between the load-on and load-off conditions, consistent with patterns observed in individual neurons (c.f. cells 1, 3, and 6 in Fig. 3B). Because inactivation of motor cortex had no effect on load compensation, this dominant, load-related signal was output-null. In addition, two smaller load-related dimensions were identified, and both were modulated by step phase, though these could potentially result from small shifts in spike alignment^46^. Because prior work has shown cortical compensation for load during voluntary upper limb movements can be influenced by ascending cerebellar drive^10,11^, we next tested whether the load signals observed during locomotion were cerebellum-dependent by examining several consequences of cerebellar perturbation. First, the effect of Purkinje cell stimulation was concentrated primarily in a single dimension (11.8% of the variance; Fig. 4C) and, like the load effect, consisted of a tonic shift in activity (Fig. 4A, center; Fig. 4B, third column). Second, the principal axes with the largest effects of load and cerebellar perturbation were not closely aligned (inner product −0.45; Fig. 4C, upper triangular matrix), though we failed to reject the null hypothesis that their relative orientations were random (p = 0.31, exact test based on beta distribution; Fig. 4C, lower triangular matrix). Third, the interaction between load and cerebellar perturbation was small, accounting for only 1.0% of firing rate variance (Fig. 4B, fifth column, third row; Fig. 4C). Fourth, activity in the top load-related dimension was not partitioned by cerebellar perturbation; instead, trajectories were tightly grouped within each load condition (Fig. 4A, left; Fig. 4B, second column, first row). Finally, projection of firing rates aligned to the onset of cerebellar perturbation onto the top load dimension revealed a minimal response (Fig. 4D, upper), while projection onto the first Purkinje cell stimulation dimension produced a large signal that was sustained throughout the stimulus train (Fig. 4D, lower). Taken together, these observations support the hypothesis that the cortical representation of load in the adaptive locomotion task is independent of cerebellar input.

### 2.5. Effects of load and cerebellar perturbation on spinal motoneuron dynamics

How are the dynamics observed in motor cortex related to the final output of the nervous system at the level of spinal motoneurons? In healthy motor units, muscle fiber action potentials are tightly locked to action potentials in the corresponding motoneuron. Thus, to address this question, we implanted flexible, high-density electrode arrays^48^ and fine wire electrodes in the forelimb muscles (see Methods), enabling us to record motor output at the resolution of individual spinally-innervated motor units in the adaptive locomotion task (Fig. 5A; n = 108 motor units, n = 27 sessions, n = 6 L7Cre-2 x Ai32 mice). Inertial load and cerebellar perturbation were applied as in the cortical recording experiments. Motor units were more strongly entrained to the locomotor rhythm (n = 108/108, q < .05, Rayleigh test) in comparison to cortical units, with flexor motor units activated during swing, and extensor motor units during stance (Fig. 5B). The firing rates of 50.9% of motor units were significantly modulated by load (n = 55/108; q < .05, multi-way ANOVA; Fig. 5C-D), 28.7% by Purkinje cell stimulation (n = 31/108), 20.4% by both load and stimulation (n = 22/108), and 7.4% by the interaction between load and stimulation (n = 8/108). Among the load-sensitive neurons, 38.2% had firing rate increases, while increases occurred in 71.0% of Purkinje cell stimulation-sensitive neurons. To identify coordinated activity patterns at the motoneuron population level, we performed dPCA as for the cortical population, and projected firing rates onto twenty dPCA decoder dimensions, which explained 94.2% of the total firing rate variance. The dominant patterns revealed by dPCA consisted of robust, condition-invariant oscillations (Fig. 6A, X-Y axes; Fig. 6B, first column), and overall, the condition-invariant signals accounted for 70.6% of explained firing rate variance (Fig. 6C). The first two condition-invariant dimensions showed approximately sinusoidal oscillations with a period of one stride, while the third had a period of one-half stride (Fig. 6B, first column). Inertial load and Purkinje cell stimulation had modest effects, accounting for 7.4% and 4.6% of the variance, respectively (Fig. 6A, left and center; Fig. 6B, second and third columns; Fig. 6C). In contrast with cortical activity patterns, the first load and Purkinje cell stimulation dimensions for the motoneuron population exhibited a clear dependence on step phase, with maximal separation between conditions in mid-stance. Animal speed accounted for 12.8% of the firing rate variance, with continuous, tonic shifts in the first speed dimensions, and more complex, step-phase-dependent effects in the second (Fig. 6B, fourth column). While these patterns yielded stacked, elliptical trajectories in the first two condition-invariant dimensions and the first speed dimension (Fig. 6A, right), roughly resembling the corresponding cortical dynamics (c.f. Fig. 4A, right), these spinal trajectories were less clearly separated across speed conditions in comparison to cortex. Finally, projection of motoneuron firing rates aligned to Purkinje cell stimulation onto the dPCA axes revealed no effect on load dimensions, but a small, tonic modulation in the first two stimulation dimensions (Fig. 6D), though these were small in comparison to the corresponding cortical signal (c.f. Fig. 4D).

**FIGURE 5.**
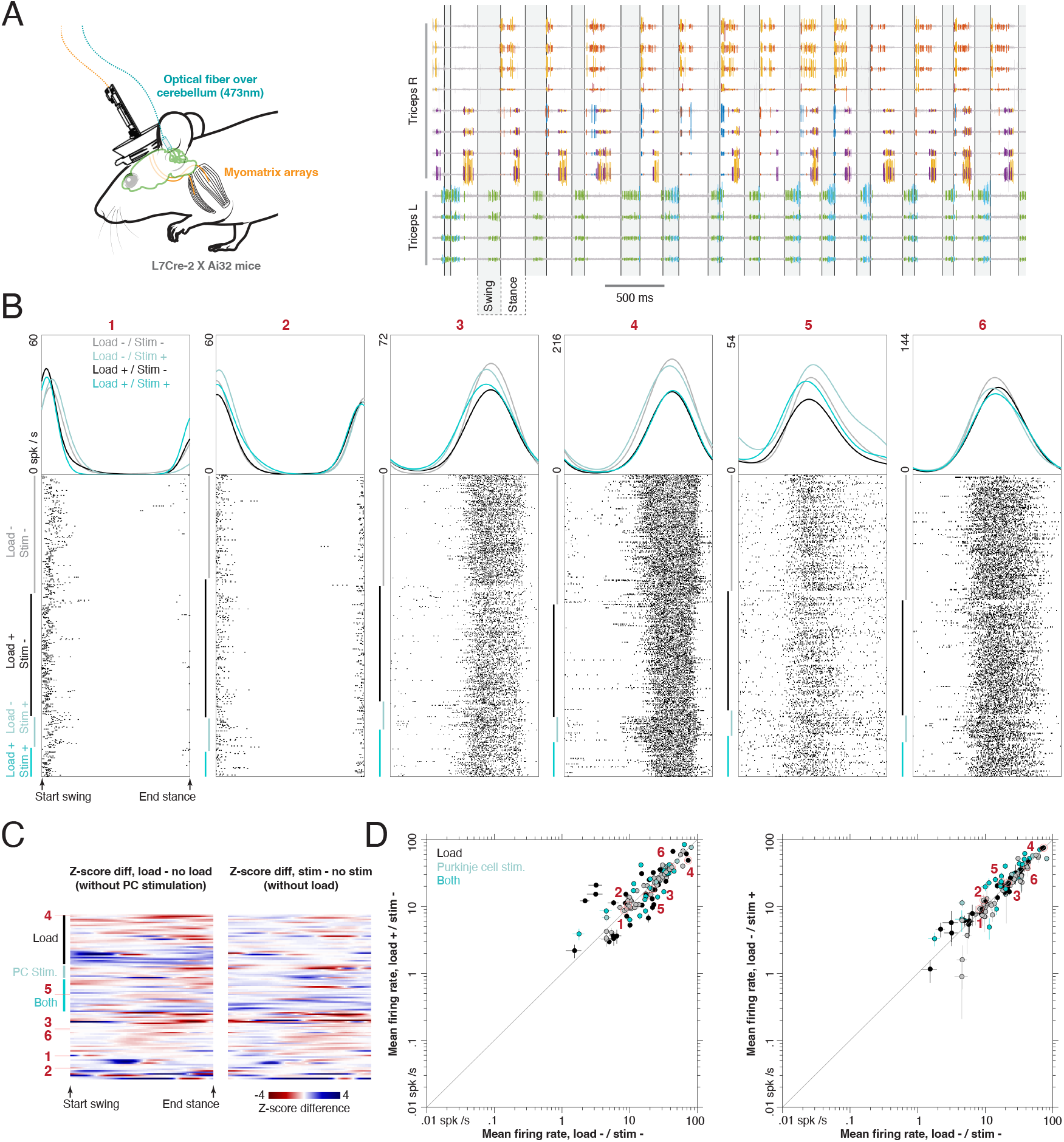
Effects of load and cerebellar perturbation on spinal motoneurons. (A) Left: mice (n = 6, L7Cre-2 x Ai32) were chronically implanted with fine wire electrodes or Myomatrix electrode arrays in the biceps brachii and triceps brachii muscles and an optical fiber over the ipsilateral cerebellar cortex for stimulation of Purkinje cells. Right: raw motor unit data recorded from the Myomatrix array during locomotion. (B) Firing rates and spike rasters for six spinally-innervated motor units in the adaptive locomotion task. Units 1 and 2 were recorded from the biceps ipsilateral to the load, and units 3-6 from the ipsilateral triceps. (C) Effect of load and cerebellar perturbation on motor units (n = 108). Each row corresponds to a single motor unit, and displays the difference in z-scored firing rate between load on and off conditions (left) and between Purkinje cell stimulation on and off (right). Units are grouped based on the detection of an effect of load (black bar), stimulation (light blue bar), both (dark blue bar), or neither (remaining neurons). (D) Mean firing rates for all motor units in load on and off conditions (left) and Purkinje cell stimulation on and off (right). Color code reflects the detection of effects of load, stimulation, or both, as in (C). Bars indicate 95% confidence intervals for the mean.

**FIGURE 6.**
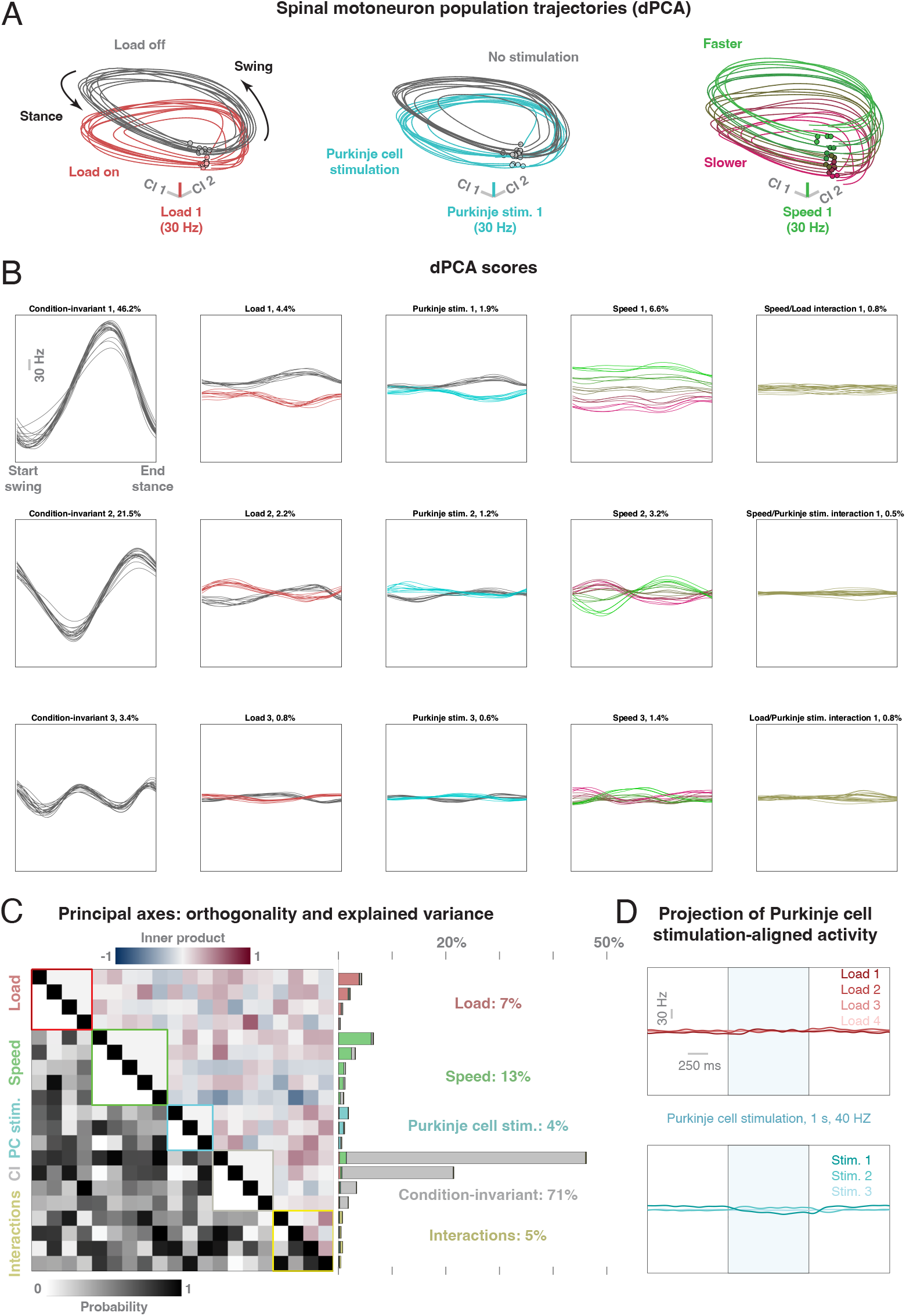
Neural population dynamics in spinal motoneurons. (A) Step-aligned neural trajectories obtained by demixed principal component analysis (dPCA). The x and y axes in each panel represent population activity in the first two condition-invariant (CI) dimensions across twenty conditions. The z axes represent the first load dimension (left), Purkinje cell stimulation dimension (center), and speed dimension (right). (B) Step-aligned neural activity across CI, load, Purkinje cell stimulation, speed, and interaction dimensions. (C) Inner product between the principal axes (left, upper triangular), probability the inner product is larger than the observed value for randomly-oriented vectors (left, lower triangular), and fraction of variance explained by each dimension (right). (D) Projection of firing rates aligned to Purkinje cell stimulation onto load dimensions (upper) and Purkinje cell stimulation dimensions (lower).

### 2.6. Distinct dynamics in cortical and spinal motoneuron populations

Although neural dynamics in the cortical and spinal populations had several qualitative similarities, including the shape of trajectories in the leading condition-invariant and speed dimensions, several key differences were apparent. First, the condition-invariant dimensions had similar time-varying trajectories (Fig 4B and 6B, first column), but the amount of firing rate variance explained was 2.4-fold larger in the spinal motoneuron population (70.6% and 28.8% in spinal and motor cortex, respectively). In this sense, the spinal population response primarily reflected the locomotor rhythm, while load, speed, and Purkinje cell stimulation imposed smaller modulations on this rhythm. In motor cortex, however, the dominant signal was related to load, and components for both Purkinje cell stimulation and speed were also prominent. This larger balance of condition-invariant activity for spinal motor output in comparison with cortex in locomoting mice contrasts with findings in primates reaching to multiple targets, which showed a much larger condition-invariant component in cortex^49^, and in primates walking over obstacles^45^. Second, load and Purkinje cell stimulation effects for motor cortex consisted primarily of tonic shifts, whereas the corresponding effects on spinal motoneurons were modulated by step phase. Third, neural trajectories were more clearly separated at different speeds for the cortical than for the spinal population.

To determine how these differences influenced the geometry of neural trajectories in the two populations, we next modeled the effects of each experimental variable by estimating maps from trajectories in one set of conditions to trajectories in a second set of conditions. In particular, for each neural population (motor cortex and spinal motoneurons) and variable (load, Purkinje cell stimulation, and speed), we used Procrustes analysis to identify the rotation, translation, and rescaling required to map trajectories in baseline conditions (load-off, stimulation-off, and lowest speed) to the corresponding trajectories in the complementary conditions (load-on, stimulation-on, and highest speed; see Methods). This produced a concise description of how each experimental manipulation altered neural trajectory geometry. Inertial load and Purkinje cell stimulation induced large vertical translation in the cortical trajectories (Fig. 7A, left and center), but produced largely rotational effects for spinal trajectories (Fig. 7B, left and center; Fig. 7C, left and center), resulting from the modulation by step phase in the latter case. For speed, both populations displayed a combination of rotation and translation, along with a slight increase in scale (Fig. 7A-C, right). We also observed that, while cortical trajectories were clearly separated across experimental conditions, spinal trajectories had greater overlap across conditions and time points. To quantify this finding, we computed a trajectory tangling index^50^ (see Methods), which measures the extent to which nearby neural states have distinct derivatives. We found that trajectories in motor cortex consistently exhibited lower tangling in comparison with the spinal motoneuron population (Fig. 7D). Highly tangled trajectories imply dynamics that are driven by external input, while low tangling may suggest more autonomous dynamics that are robust to noise. However, motor cortex maintains relatively low tangling despite the presence of strong signals about the state of the limbs and throughout experimental manipulation of cerebellar inputs. Thus, low tangling might constitute a mark of noise robustness even in systems that depend strongly on inputs.

**FIGURE 7.**
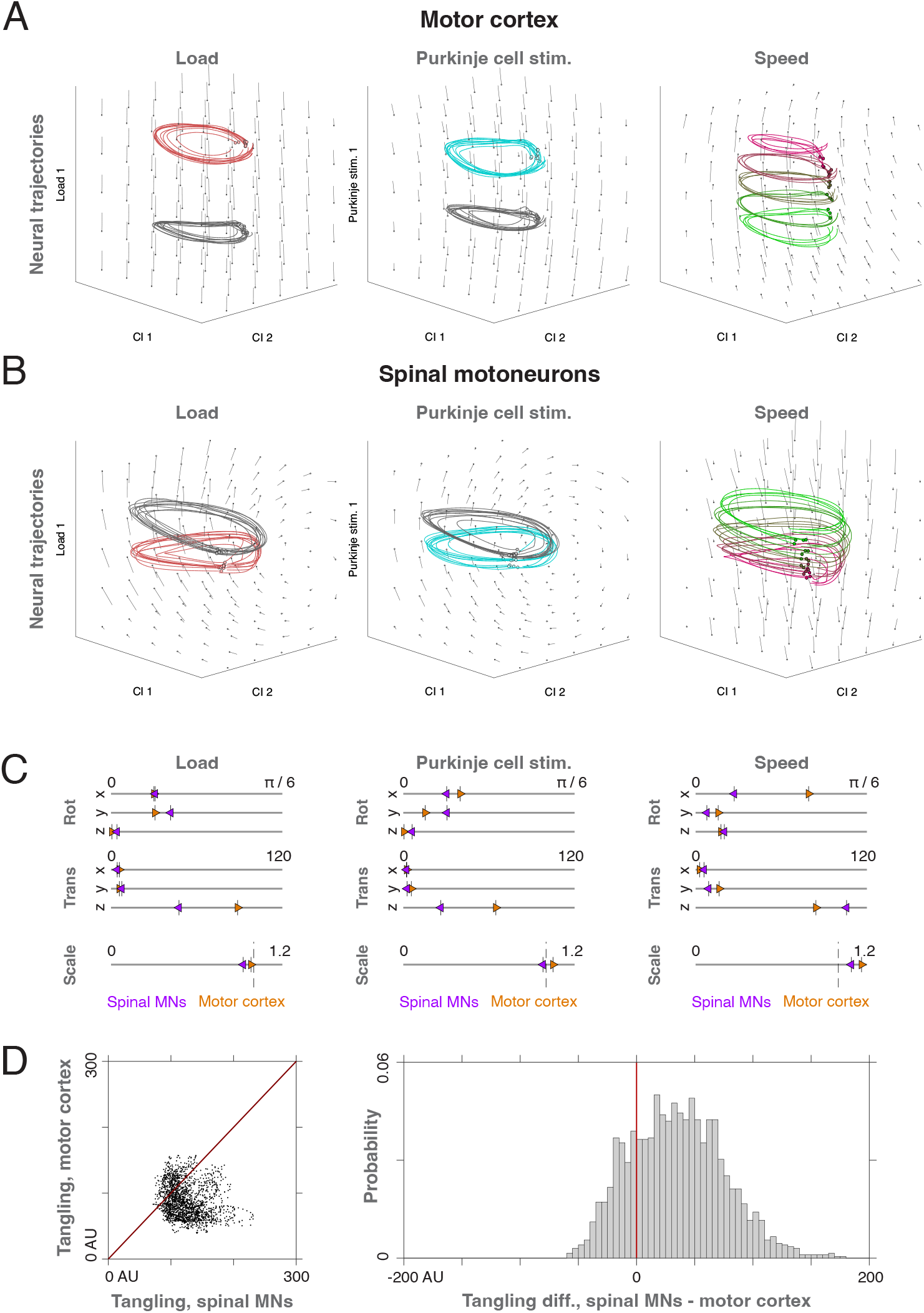
Comparison of neural population geometry in motor cortex and spinal motoneurons. (A) Neural trajectories and Procrustes transformations for the motor cortex population. In each panel, the vector field indicates the direction and magnitude of the Procrustes transformation required to map the trajectories from one set of conditions onto the trajectories for another set. Left: map from load-off trajectories to load-on trajectories. Center: map from stimulation-off trajectories to stimulation-on trajectories. Right: map from slowest trajectories to fastest trajectories. (B) Neural trajectories and Procrustes transformations for the spinal motoneuron population. Conventions as in (A). (C) Comparison of Procrustes transformations for the motoneuronal and cortical populations. (D) Comparison of trajectory tangling in motoneurons and motor cortex. Left: scatterplot of tangling values for motoneurons and cortex across all conditions and time bins. Right: histogram of differences in tangling between motoneurons and cortex.

## 3. Discussion

In this study, we have identified a robust signature of limb inertial load in the mouse motor cortex during adaptive locomotion, which comprised the largest component of cortical activity in the task. Because muscle activity during load compensation was unchanged by cortical inactivation, we conclude this load-related signal is not a motor command underlying the compensation, but is instead a latent, output-null representation. Our finding that activity along the load dimension is minimally influenced by cerebellar perturbation further suggests it is not driven primarily by cerebellar projections through the ventrolateral thalamus, but may reflect sensory signals ascending from the dorsal column nuclei via somatosensory cortex^51^ or a more abstract contextual signal from other cortical regions. This output-null representation of load may support the generation of appropriately-scaled commands when a voluntary, cortically-dependent gait modification must be integrated with the spinally-generated locomotor program. For example, during locomotion over obstacles, which requires motor cortex in cats^29–31^ and mice^33^, an animal must generate larger flexor torques at higher loads, and a latent change in activity along load-related dimensions might increase the amplitude of a cortical command for voluntary adaptation of gait. In addition, the load-related signals we observe in cortex might modulate spinal reflex gains to adjust the motor response to unexpected perturbations, as has been found for rhythmic, voluntary upper limb movements^52^ and during split-belt locomotor adaptation in humans^53^.

The motor cortical dynamics we observed share several key similarities with those reported in primates performing a voluntary cycling task^47,50,54^. Neural trajectories in primary and dorsal premotor cortex during cycling are periodic and elliptical in the dominant dimensions, and translate continuously along an axis approximately orthogonal to the plane of rotation with changing speed. These dynamics are consistent with a cortical rhythm generator that determines movement speed and phase while driving smaller, more complex, muscle-like output commands that control movement via corticospinal projections. In locomotion, by contrast, the rhythm is generated by an intrinsic spinal circuit, and oscillatory activity in the cortical condition-invariant dimensions likely reflects sensory feedback or an efference copy from the CPG. Thus, although the condition-invariant activity in mouse spinal motoneurons closely resembles these cortical dynamics, it is unlikely they are driven by cortical commands. Indeed, we observed that inactivation of motor cortex had little effect on either the rhythmic flexor-extensor alternation or on the additional forelimb EMG changes imposed by load. Another feature of primate cortical dynamics during cycling is the maintenance of significantly lower trajectory tangling in comparison with muscle activity. That is, nearby neural states have similar derivatives, so cortical trajectories tend to avoid crossing one another across different time points and conditions. Because higher tangling is a signature of external forcing, low tangling is consistent with strong internal dynamics in the primate cortical network during the task. In locomoting mice, we also observe lower levels of tangling in motor cortex in comparison to the spinal motoneuron population, which must be driven by external inputs. This difference, however, is smaller than in the primate cycling task, consistent with a spinal rather than cortical locus of pattern generation, and with a greater role for inputs in driving cortical dynamics. In addition, cycling studies used both forward and backward rotations, which tended to increase tangling in muscle trajectories, while we tested locomotion in the forward direction only.

Our findings highlight a disassociation between the dominant patterns of motor cortical activity in a given task and the necessity of these patterns for generating motor output. Because many distinct descending and spinal pathways ultimately converge onto the same motoneurons, the problem of inferring the effects of cortical dynamics on muscle activity from simultaneous measurements of both is necessarily ill-posed. Furthermore, changes in cortical activity with experimental conditions or behavioral epoch may effectively cancel out at the motoneuronal level, enabling cortical computations to occur without influencing movement^21,22^. An emerging body of evidence suggests the contribution of motor cortex to forelimb movements can depend strongly on behavioral task and context. In the mouse, silencing motor cortex has negligible effects on normal locomotion^32,33^, moderately impairs skilled gait modification^33^, and severely disrupts precise reach-to-grasp movements^20,55,56^. Correlations between cortical neurons and the mapping between neural and muscle activity can change substantially between tasks^32,57^, though work in the cat suggests this mapping is preserved between voluntary gait modification and reaching^58^. In rats, lesions to motor cortex impair learning of an interval timing task, but do not affect performance if delivered after the task has been learned^59^, and the necessity of motor cortex for a task can depend on the preceding training regimen^60^. Meanwhile, studies of neural population dynamics in reaching primates have emphasized the significance of cortical dimensions that are decoupled from movement and contribute to internal computations during motor preparation^21,22^, initiation^49,61^, and learning^62,63^. Our results build upon these findings by identifying a robust, latent representation of limb mechanics in motor cortical population activity during the adaptation of a rhythmic movement governed by a spinal CPG.

## 4. Methods

### 4.1. Experimental animals and behavioral task

All experiments and procedures were approved by the Institutional Animal Care and Use Committee at Case Western Reserve University, and in accordance with NIH guidelines. At the time of surgical implantation, mice were 16-23 weeks old and weighed approximately 24-33 g. Mice with higher body mass were selected for experiments, as they were better able to carry the implant payload on the head. A total of 12 adult mice were used for experiments, including four (male) VGAT-ChR2-EYFP line-8 strain mice (Jackson Laboratory) and eight (six male and two female) L7Cre-2 x Ai32 strain mice (Jackson Laboratory). Animals were healthy, individually housed under a 12-hour light-dark cycle, and had no prior treatment, drug or altered diet exposure. After surgery, animals were cared for and studied for up to three months.

### 4.2. General surgical procedures

All mice were implanted with optical fibers for optogenetic perturbation, and with either (1) fine wire electrodes in forelimb muscles for electromyographic (EMG) recording, (2) Myomatrix arrays^48^ for high-resolution recording from motor units, or (3) silicon probes in motor cortex for neural ensemble recording. The initial surgical procedures preceding implantation of EMG or neural electrodes was similar across surgeries. Anesthesia was induced with isoflurane (1-5%, Kent Scientific), eye lubricant was applied, fur on top of the head and posterior neck was shaved, and the mouse was positioned in a stereotaxic apparatus (model 1900, KOPF instruments) on top of a heating pad.

Under sterile technique, the top of the head was cleansed with alternating swabs of 70% ethanol and iodine surgical scrub, lidocaine (10 mg/kg) was injected under the skin on the top of the skull, the skin was removed, the periosteum on top of the skull removed, and a custom designed 3D-printed head post was attached with UV-cured dental cement (3M RelyX Unicem 2). Then, optical fibers and chronic recording electrodes were surgically implanted (see below). Post-surgery, the minimum recovery period was 48 hours, Meloxicam (5 mg/kg) was administered for pain management once per day, and the investigators monitored animal behavior, body mass, food and water intake on a daily basis. The recovery period was extended an additional 24-48 hours for some animals as necessary.

### 4.3. Adaptive locomotion task

After at least two days of recovery from surgery, mice were placed on a custom-built motor-driven treadmill (46 cm long by 8 cm wide) that was controlled at fixed speeds between 10-30 cm/s (Fig. 1B). The treadmill apparatus was enclosed in transparent acrylic, and belt speed monitored by a rotary encoder. Locomotion was motivated through negative reinforcement with airpuffs triggered by an infrared brake beam at the back of the treadmill belt. Mice were acclimated to the apparatus for up to three sessions, until they ran continuously without prompting. For the condition of unrestrained, load adaptive locomotion, one investigator briefly scruffed the mouse while another positioned a small weight (0.5 g) on the wrist, and at the conclusion of the load-on condition the wrist weight was removed. The wrist weight was fabricated by gluing a steel ball bearing to a small zip-tie. For each animal, recording sessions were performed up to twice a day. Per session, mice ran between 5-20 min within the load-off and 5-10 min within the load-on conditions. Sessions started with the mouse running in the load-off condition that was followed by load-on, in a subset of sessions (n = 8) there was a final load-off condition that was performed. Each session was concluded based on mouse performance having at least 5 min of continuous locomotion per condition, or was ended due to mouse stress or reaching the 30 min time mark.

### 4.4. Videography

Four synchronized high speed cameras (Blackfly, model BFS-U3-16S2C-CS, Teledyne FLIR; Vari-Focal IP/CCTV lens, model 12VM412ASIR, Tamron) were positioned around the treadmill, with two cameras recording from each side of the treadmill belt, acquiring approximately sagittal views of the locomoting mouse. Under infrared illumination of the field, each camera was positioned to record the complete length of the treadmill belt at a frame rate of 150 Hz and a region of interest of 1440 x 210 pixels, and was triggered by an external pulse generator using custom LabVIEW code (National Instruments). Images were acquired with the SpinView GUI (Spinnaker SDK software, Teledyne FLIR).

### 4.5. Pose estimation during locomotion

For tracking mouse pose (i.e., anatomical landmarks) across cameras during locomotion, DeepLabCut^37^ was used. The position of 22 landmarks was tracked, including the nose, eye, fingertip, wrist, elbow, shoulder, toe, foot, ankle, knee, hip, and tail on each side of the body. Separate tracking models were developed for EMG and cortical electrodes due to differences in animal appearance between the implant types. In total, 1850 and 2002 labeled frames were used for training the EMG and cortical implant models, respectively. Next, Anipose^38^ was used to triangulate 3D pose from the 2D estimates in the four cameras. Briefly, the four cameras were calibrated using simultaneously-acquired images of a ChArUco board, and the 3D pose estimated by minimizing an objective that enforced small reprojection errors, temporal smoothness, and soft constraints on the length of rigid body segments.

The pose estimates obtained from Anipose were then transformed into a natural coordinate frame: (1) forward on treadmill, (2) right on treadmill, and (3) upward against gravity. Next, the forward coordinate was unrolled by adding the cumulative displacement of the treadmill computed from the rotary encoder. This resulted in a treadmill-belt-centered coordinate frame, as though the mouse was progressing along an infinitely-long track: (1) forward on treadmill, relative to the unrolled position of the back of the belt at the start of the experiment, (2) right on treadmill, and (3) upward against gravity. Sessions were then segmented into swing and stance epochs by detecting threshold crossings of the forward finger velocity and upward finger position. The identified swing and stance time points were used for alignment of electrophysiological recordings. For each mouse and session, the quality of the pose estimates was assessed using Anipose quality metrics, visual inspection of trajectories, and comparison of estimated pose with the raw videos.

### 4.6. Optogenetic perturbations

Optical fibers (catalog number FT200UMT, fiber core diameter 200 µm, ThorLabs) were glued inside ceramic ferrules (catalog number CFLC230-10, ThorLabs) and positioned onto the skull over a thin layer of transparent dental cement (Optibond, Kerr), which enabled optical access to the brain^43,64^. Ferrules were placed bilaterally above the forelimb motor cortex (bregma +0.5 mm, lateral 1.7 mm) of VGAT-ChR2-EYFP mice to stimulate inhibitory interneurons^20,32,65^. Separately, ferrules were placed bilaterally above the pars intermedia of cerebellar lobule V (bregma −6.75 mm, lateral 1.7 mm) of L7Cre-2 x Ai32 mice to stimulate Purkinje cells^66,67^.

Optogenetic perturbation with a 473 nm wavelength laser was delivered with sinusoidal waves at 40 Hz (Opto Engine LLC). The laser was triggered by an external signal generator controlled with custom labVIEW software. Power levels used during locomotion were based on average ranges from prior investigations^20,32,42,66,67^. In L7-Cre-2 x Ai32 mice, optogenetic perturbation of the cerebellum at higher power levels stopped mouse locomotion, and the forelimb musculature was unable to support the mouse during stance. We therefore adjusted the power level for each animal based on the effects of stimulation in the home cage.

Home cage sessions (Supplemental Fig. 1A) involved stepwise power level adjustments of the optogenetic perturbation and measurement of EMG. In L7-Cre-2 x Ai32 mice, we found that Purkinje cell stimulation (0.125-4 mW) induced suppression of forelimb flexor and extensor EMG, followed by a rebound response after the termination of the stimulus. We therefore adjusted the laser power for behavioral sessions to a level that produced minimal rebound, and did not halt locomotion (0.25-2 mW). Likewise, stepwise power level adjustments were made to confirm quiescent muscle activity in VGAT-ChR2-EYFP mice (1-12 mW), and power levels at the high end of this range (8-12 mW) were then used for behavioral experiments. For home cage sessions, the stimulus was a 40 Hz sine wave with a duration of 0.25, 0.5 or 1 s, and interstimulus intervals were randomized and between 3-10 seconds.

### 4.7. Electromyogram recordings

Electromyogram (EMG) recordings of gross muscle activity from the elbow flexors and extensors was made using fine-wire^32,68,69^ electrodes, and recordings from single motor units were performed with both fine-wire electrodes and high-density Myomatrix arrays^48,70,71^. For each mouse, we implanted a total of four muscle locations, targeting an elbow flexor and extensor muscle on each side. Fine-wire electrodes were made with four pairs of wires in a bipolar EMG configuration, following an established protocol^68^. Each bipolar fine-wire electrode comprised two 0.001 inch diameter, seven-stranded braided steel wires (catalog number: 793200, A-M Systems) that were crimped into a 27 gauge needle, twisted and knotted together. For recording contacts, ∼0.5-1 mm of insulation was removed per wire between the knot and needle, made closer to the knot, and staggered with an inter-contact distance of ∼2 mm. The open ends of the wire on the other side of the knot were soldered onto a 32-pin connector (Omnetics Nano, A79025, 36 pins, 4 guideposts), along with a gold pin cap for attachment to the ground (Mcmaster-Carr). Myomatrix electrodes^48^ were used to only record EMG with single motor unit resolution, these electrodes had gold contacts that were plated with conductive polymer PEDOT to reduce the impedance to the measured range of 3-23 kΩ. Fine-wire electrodes were grounded with a gold pin soldered to a stainless steel wire placed through the skull and into the brain by performing a craniotomy with a dental drill ∼4 mm rostral to the forelimb area of motor cortex area. The dura was left intact, Kwik-sil (World Precision Instruments) was applied, and the pin was secured to the skull with dental cement. Myomatrix electrodes were grounded onto the skull and secured with dental cement.

For surgical implantation, the fur on the posterior neck, posterior shoulders and both forelimbs above the elbow joint of the mouse was removed using depilatory cream prior to positioning within the stereotaxic apparatus. Electrodes were implanted only after the headpost, optical fibers and ground were secured. For each forelimb, lidocaine was injected under the skin, and a 2-3 cm incision of the skin was made between the elbow and shoulder joint, along the midline axis of the lateral head of the triceps brachii muscle, and was subsequently kept moist with saline. Each electrode was led under the skin from the posterior neck to be separately implanted in the long head of the biceps brachii or triceps brachii muscles. For targeting the biceps brachii muscle the forelimb was abducted, elbow extended and the paw supinated, whereas for targeting the triceps brachii muscle, the elbow was flexed and paw pronated. The skin was adjusted using forceps to provide an opening over the targeted muscle, and electrodes were inserted into the muscle belly from proximal to distal. The fine-wire electrodes were inserted with the attached crimped needle, after insertion, the needle and excess distal wire was cut and a distal knot was made. For Myomatrix electrodes, a suture knot was tied onto the distal polyimide hole of each thread, then, following the suture needle, was carefully pulled into the targeted muscle belly. One Myomatrix thread was inserted per muscle. For both the fine-wire and Myomatrix electrode implants, the incised skin was then flushed with saline and sutured. The connector was then secured to the head post with dental cement and the inferior skin relative to the head post was hermetically sealed with skin adhesive (3M Vetbond).

Despite targeting muscle long heads during implantations, we did not systematically differentiate EMG between the long and short head of the biceps brachii muscle, and likely EMG during locomotor swing comprised the synergist contribution from other elbow flexor muscles including the brachialis and coracobrachialis. Likewise, we did not differentiate EMG between the heads of the triceps brachii muscle, and it remains possible that EMG during stance may have had minor synergist contribution from the dorso-epitrochlearis brachii and anconeus muscles^72^.

We implanted EMG electrodes in forelimb muscles bilaterally, because throughout the course of experiments the signal-to-noise would degrade and in some instances electrodes would be damaged, and these sessions were excluded. Therefore, the forelimb with better EMG signal-to-noise and minimal crosstalk from other muscles was used for experiments, determining on which side the wrist weight and optogenetic perturbations were applied. In VGAT-ChR2-EYFP mice, optogenetic silencing of the forelimb area of motor cortex was linked to contralateral forelimb EMG and contralateral load. In L7-Cre-2 x Ai32 mice, optogenetic silencing of deep cerebellar nuclei through the activation Purkinje cells was linked to ipsilateral forelimb EMG, ipsilateral load, and contralateral cortical neuron recordings. Three (one female) L7-Cre-2 x Ai32 and four VGAT-ChR2-EYFP mice were implanted with fine-wire electrodes and three (one female) L7-Cre-2 x Ai32 mice were implanted with Myomatrix electrodes. Recordings were amplified and bandpass filtered (0.01-10 kHz) using a differential amplifier and digitized (Intan RHD2216, 16-bit, 16 channel bipolar input recording headstage), and acquired at 30 kHz (Open Ephys acquisition board and software). At the conclusion of experiments on each mouse, the targeted muscles were verified post-euthanasia by dissection.

For subsequent analysis of step-aligned muscle activity, the gross EMG was high-pass filtered (200-250 Hz cutoff), rectified, and convolved with a Gaussian kernel (σ = 10 ms). To normalize the smoothed EMG signal, we first detected all peak events exceeding the 90th percentile of the full time series. Then, the smoothed signal was divided by the median amplitude of these peaks.

### 4.8. Motor unit spike sorting

On many EMG recordings from fine-wire electrodes, single motor units were identified (e.g., the triceps unit in Fig. 1C-D). For these fine-wire recordings, the EMG was high-pass filtered on each channel (cutoff set between 200 and 1000 Hz, 2nd order Butterworth). Motor unit spike times were identified by voltage threshold and waveform template matching (Spike2 software, version 7, Cambridge Electronics Design). In the fine-wire electrodes implanted in the biceps brachii muscle, single motor units were sometimes recorded during the stance phase, possibly due to the small relative volume of elbow flexor to extensor muscle and that the cut-end of the electrode was closer to the distal aspect of the lateral triceps brachii.

For Myomatrix electrodes, each thread comprised four bipolar recording channels that were implanted into the same muscle that enabled correlated voltage and waveform analysis across channels. The EMG was high pass filtered (400-500 Hz cutoff, Parks-Mclellan method), and motor unit waveforms and spike times were extracted using an existing method^73^. Then, clusters were manually cut using peak-to-trough features from all channels on each thread, and unit quality assessed by inspection of waveforms, autocorrelations, cross correlations between units recorded on the same thread, and raw signals with unit spike times superimposed. Overall, we recorded 54 ipsilateral extensor units, 27 contralateral extensor units, 17 ipsilateral flexor units, and 10 contralateral flexor units.

### 4.9. Motor cortical recordings

Extracellular recordings in the forelimb area of the motor cortex^20,74^ were made using chronically implanted high-density silicon probes (64 channel, 4-shank, 6 mm length E1 probe, Cambridge NeuroTech) secured to a manual micromanipulator (CN-01 V1, Cambridge NeuroTech). Probes were plated with the conductive polymer PEDOT to reduce the impedance to the measured range of 30-50 kΩ, and the tips were sharpened to ease insertion through the dura. The electrode was grounded with a gold pin soldered to a stainless steel wire placed through the skull and into the visual cortex. Surgical implantation of the probe occurred after the headpost, optical fibers and ground were secured to the skull. A craniotomy (dimensions ∼1×2 mm) was performed with a dental drill to access the forelimb area of motor cortex on the left side (bregma +0.5 mm, lateral 1.7 mm), care was taken to leave the dura intact, and cold saline was applied continuously to reduce swelling. The probe tip was inserted to a starting depth between 400-540 µm, silicone gel was applied (catalog number 3-4680, Dowsil, Dow) and the apparatus including the amplifier was secured to the head post, skull and enclosed custom chamber using dental cement.

Two L7-Cre-2 x Ai32 mice were implanted and recordings were amplified and band-pass filtered (0.01-10 kHz) using a differential amplifier and digitized (mini-amp-64, Cambridge NeuroTech) and acquired at 30 kHz (Open Ephys GUI). Each session, the electrophysiological signal-to-noise and spiking density across channels was assessed, to record from new neurons and when signal quality degraded, the probe was moved 62.5-125 µm deeper every 1-3 days by adjusting the micromanipulator until the lowest recording channel hit white matter (∼1-1.2 mm from the surface).

### 4.10. Motor cortex spike sorting

Single units in the motor cortex were identified using Kilosort 2.5^75–77^ (https://github.com/MouseLand/Kilosort), and manually curated with the Phy GUI (https://github.com/cortex-lab/phy). Only well-isolated neurons were accepted based on spike waveforms, the presence of an absolute refractory period greater than 1 ms, the stability of spike amplitude over the session, and isolation of the cluster in feature space. Spike time cross-correlation was used to remove duplicated neurons.

### 4.11. EMG analysis

To assess changes in behavior over individual experimental sessions, we first interpolated the smoothed biceps and triceps EMG and forward finger velocity between the start of swing and end of stance on each step cycle, and visualized the resulting curves as heatmaps (Fig. 1E). For optogenetic perturbation experiments, we averaged the step-aligned curves within each condition (load on / off, optogenetic perturbation on / off), and visualized the means using polar plots (Fig. 2A-B). Next, to obtain a compact representation of motor output on each step, we averaged the biceps (flexor) EMG during the swing epoch, the triceps (extensor) EMG during the stance epoch, and velocity (fingertip) over the entire step. Medians and bootstrapped confidence intervals for load-off and load-on conditions were visualized as scatterplots (Fig. 1F), and a difference between conditions (where each paired observation is a load-off and load-on median in one session) assessed with a two-sided sign rank test. The trend in step-averaged EMG across each session was modeled using loess smoothing^78^ (second-order, smoothing parameter α = .9; Fig. 1G). To estimate the effects of load, optogenetic perturbation, and speed on EMG and velocity, we fit one linear model for each session using ordinary least squares, where each observation corresponded to a single step. The dependent variables were biceps EMG, triceps EMG, and forward velocity, and the independent variables were step frequency (i.e., the inverse of the duration of each step), load, optogenetic perturbation, and interaction between the load and optogenetic perturbation. All variables were Z-scored to facilitate comparison of effect sizes across variables and sessions. Coefficients and 95% confidence intervals were visualized using scatterplots and histograms (Fig. 2C; Supplementary Fig. 1C), and the sign of the coefficients assessed with a sign rank test with Benjamini-Hochberg correction (q < .05; Supplementary Fig. 1D). For coefficients related to optogenetic perturbation and its interaction with load, this test was applied separately to sessions using VGAT-ChR2-EYFP and L7Cre-2 x Ai32 mice.

### 4.12. Analysis of cortical neurons and spinal motoneurons

For each motor cortical neuron and spinally-innervated motor unit, firing rates over the full experimental session were computed using Gaussian smoothing (σ = 25 ms). Using the step cycle segmentation from kinematic data (described above), smoothed firing rate curves were extracted for each step using linear interpolation between the start of swing and end of stance, then averaged within each experimental condition to create peri-event time histograms (Fig. 3B; Fig. 5B). The effects of load and Purkinje cell stimulation as a function of step phase were visualized by subtracting the Z-scored firing rates in the load-off, stim-off condition from the Z-scored firing rates in the load-on, stim-off (Fig. 3C), and load-off, stim-on conditions (Fig. 5C), respectively. Step-averaged firing rates were computed for each step by dividing the number of spikes by the step duration. Means and 95% confidence intervals for step-averaged rates were visualized with scatterplots (Fig. 3D; Fig. 5D) and analyzed with a multi-way ANOVA for each neuron. A Benjamini-Hochberg correction for multiple comparisons across neurons was applied.

### 4.13. Demixed principal component analysis

To identify the coordinated, low-dimensional dynamics in the motor cortical and spinal motoneuron populations, we used demixed principal component analysis (dPCA)^46^, which decomposes measured firing rates into latent variables related to experimental parameters of interest, using a published Matlab package (https://github.com/machenslab/dPCA). Briefly, the average step-aligned firing rate for each unit (n = 710 for cortical neurons, n = 108 for spinal motoneurons) was measured in twenty different conditions in a factorial design: load on / off (two levels) x Purkinje cell stimulation on / off (two levels) x animal speed (five levels). Firing rate was sampled at 100 evenly-spaced points across the step cycle, from the start of swing to the end of stance. For the speed factor, the forward speed of the animal’s nose at swing onset was partitioned into five bins with approximately 50% overlap using an equal count algorithm^78^. This imposed the following marginalizations over parameters: (1) load, (2) speed, (3) Purkinje cell stimulation, (4) condition-invariant, (5) load / speed interaction, (6) load / Purkinje cell stimulation interaction, and (7) speed / Purkinje cell stimulation interaction. Next, we estimated the decoder and encoder matrices with twenty components and regularization parameter λ = 1e-5, and projected firing rates onto the decoder columns to obtain scores parameterized by step phase (Fig. 4A-B; Fig. 6A-B). The alignment between pairs of principal axes was assessed by computing the inner product (Fig. 4C, upper triangular; Fig. 6C, upper triangular), and by applying an exact test against the null hypothesis that the relative orientation of the axes is random with an alternative hypothesis that the axes are orthogonal. Under the null hypothesis, (x-1)/2 follows a beta distribution with α = β = (*d* – 1)/2, where x is the inner product between axes and d = 20 is the dimension of the latent variable space. The probability the inner product x is within r of zero (i.e., that the axes are nearly orthogonal) under the null hypothesis is given by 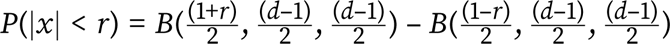, where B is the beta cumulative distribution function. Thus, setting r as the absolute value of the measured inner product between two principal axes, we can calculate the probabilities shown in Fig. 4C and 6C (lower triangular).

### 4.14. Comparison of cortical neuron and spinal motoneuron trajectories

For each neural population (motor cortex and spinal motoneuron) and experimental parameter (load, Purkinje cell stimulation, and speed), we extracted neural trajectories in the leading component corresponding to that parameter and in the first two condition-invariant dimensions across all twenty conditions. We then used Procrustes analysis within each neural population and parameter to find the optimal transformations from trajectories in one set of conditions to those in another set. These mappings could include translation, rotation, and isotropic rescaling, but not reflection. For the load and Purkinje cell stimulation parameters, trajectories in load-off and stimulation-off conditions were mapped to the corresponding trajectories in load-on and stimulation-on conditions, respectively. For the speed parameter, trajectories in the lowest speed condition were mapped to trajectories in the highest speed condition. The resulting maps were visualized on a regular 3D grid by mapping each grid point to a second point in the direction of its image under the Procrustes transformation, with a scaling of 0.2 for motor cortex and 0.4 for spinal motoneurons (Fig. 7A-B). The analysis of trajectory tangling was performed as described in previous studies^50^. Briefly, neural trajectories in the full 20-dimensional latent variable space identified by dPCA were numerically differentiated along the time axis. Next, for each time point t* and condition c*, the following quantity was computed: 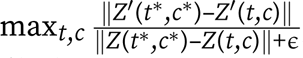, where Z(t,c) is the neural state in condition c at time t, and Z’(t,c) its derivative. The value of ɛ was set at 10% of the mean of the sum of squares of Z(t,c), concatenated across all conditions. This normalization was performed separately for the cortical and spinal populations.

## 5. Author contributions and acknowledgements

EK and BS designed the study. EK performed the experiments. KH assisted with behavioral procedures and designed the cortical recording chamber. EK and BS analyzed the data, interpreted the results, and wrote the paper. SS contributed the Myomatrix electrode arrays. BS supervised the project. We thank Andrew Pruszynski and Jonathan Michaels for discussions, Case Western Reserve University High-Performance Computing for support, and Alex Sohn for the design of the motorized treadmill. Line drawings of the mouse brain and body were provided by SciDraw.io. Research reported in this publication was supported by Case Western Reserve University, by the National Institute of Neurological Disorders and Stroke of the National Institutes of Health under award numbers R01NS129576 (PI: BS), R01NS109237 (PI: SS), R01NS084844 (PI: SS), R01EB022872 (PI: SS) and U24NS126936 (PI: SS), and by the McKnight Foundation (PI: SS), Kavli Foundation (PI: SS), Azrieli Foundation (PI: SS), and the Simons-Emory International Consortium on Motor Control (PI: SS). The content is solely the responsibility of the authors and does not necessarily represent the official views of the National Institutes of Health. EK was supported by a Postdoctoral Fellowship from the Natural Sciences and Engineering Research Council of Canada.

The authors declare no competing interests.

---

**Supplemental Figure 1.**
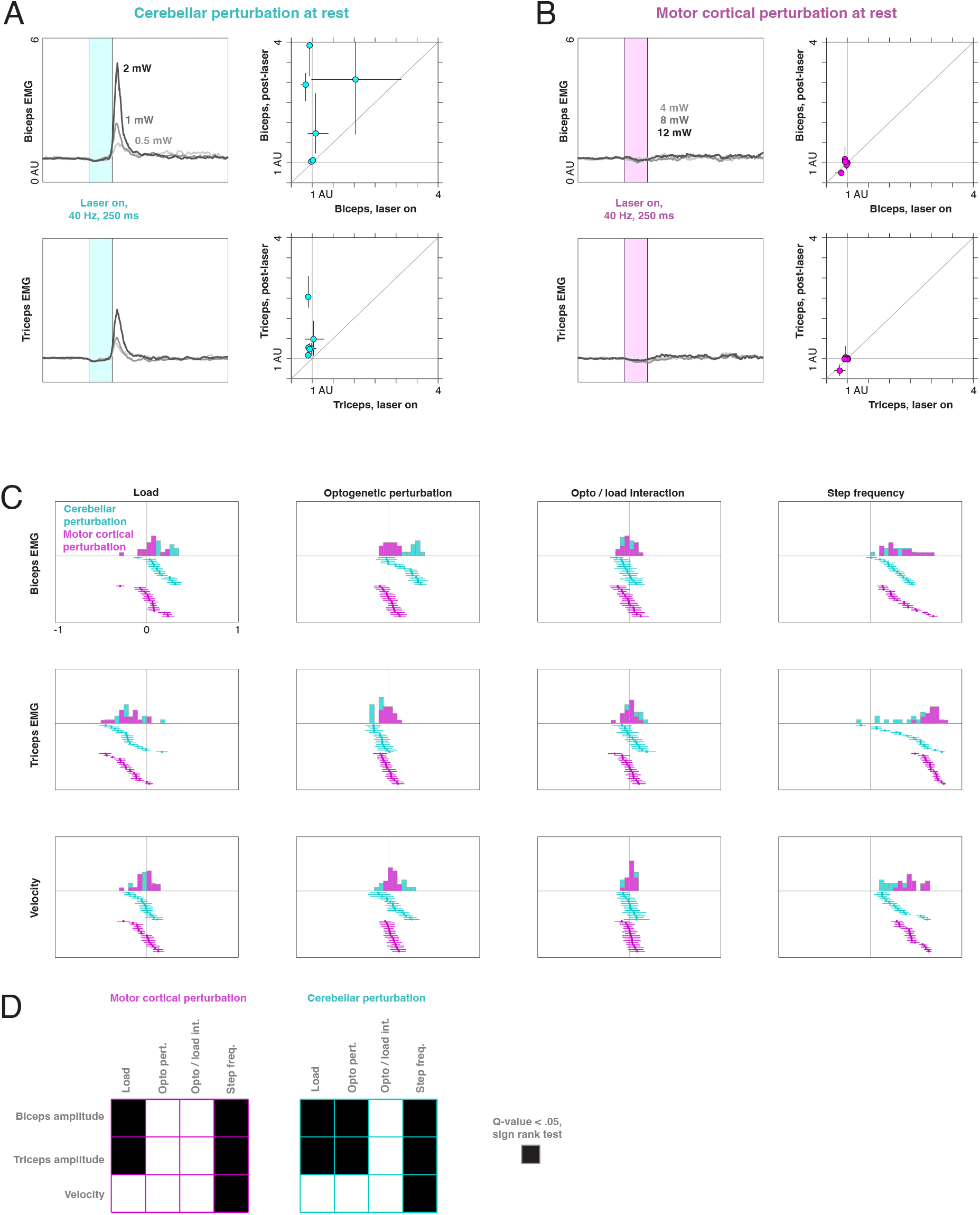
, related to Figure 2: Effects of optogenetic perturbations at rest and during locomotion. (A) Biceps and triceps EMG aligned to stimulation of contralateral Purkinje cells (40 Hz, 250 ms) in L7Cre-2 x Ai32 mice during quiet rest (n = 6 sessions, n = 3 mice). Left panels: example responses from a single experimental session. Right panels: median and bootstrapped 95% confidence intervals of EMG responses in the 100 ms following the end of stimulation, versus the responses during the stimulation epoch. Responses are normalized by the median baseline EMG preceding stimulation onset (−1000 ms to −100 ms relative to stimulation). Plotted values correspond to the highest laser power level tested in each session. (B) EMG responses to stimulation of inhibitory interneurons in the contralateral motor cortex of quietly resting VGAT-ChR2-EYFP mice (n = 6 sessions, n = 4 mice). Conventions as in (A). (C) Effects of load and optogenetic perturbation during locomotion (VGAT-ChR2-EYFP: n = 18 sessions, n = 4 mice; L7Cre-2 x Ai32: n = 16 sessions, n = 3 mice). Coefficients and 95% confidence intervals for regression of biceps EMG, triceps EMG, and forward finger velocity on load, optogenetic perturbation, the interaction between load and optogenetic perturbation, and step frequency. Each coefficient corresponds to one experimental session. (D) Outcome of one-sample, two-sided sign rank test against the null hypothesis that coefficients have median zero. Dark squares indicate q < .05 following Benjamini-Hochberg correction for multiple comparisons. Coefficients for load and step frequency were pooled between the two mouse strains.

